# MultiGreen: A multiplexing architecture for GreenGate cloning

**DOI:** 10.1101/2024.06.11.598430

**Authors:** Vincent J. Pennetti, Peter R. LaFayette, Wayne Allen Parrott

**Affiliations:** Institute of Plant Breeding, Genetics and Genomics, University of Georgia, University of Georgia, Athens, GA (USA; Center for Applied Genetic Technologies, University of Georgia, Athens, GA (USA); Department of Crop and Soil Sciences, University of Georgia, Athens, GA (USA)

## Abstract

Genetic modification of plants fundamentally relies upon customized vector designs. The ever-increasing complexity of transgenic constructs has led to increased adoption of modular cloning systems for their ease of use, cost effectiveness, and rapid prototyping. GreenGate is a modular cloning system catered specifically to designing bespoke, single transcriptional unit vectors for plant transformation— which is also its greatest flaw. MultiGreen seeks to address GreenGate’s limitations while maintaining the syntax of the original GreenGate kit. The primary limitations MultiGreen addresses are 1) multiplexing in series, 2) multiplexing in parallel, and 3) repeated cycling of transcriptional unit assembly through binary intermediates. MultiGreen efficiently concatenates bespoke transcriptional units using an additional suite of level 1acceptor vectors which serve as an assembly point for individual transcriptional units prior to final, level 2, condensation of multiple transcriptional units. Assembly with MultiGreen level 1 vectors scales at a maximal rate of 2*⌈*log*_6_*n*⌉+3 days per assembly, where *n* represents the number of transcriptional units. Further, MultiGreen level 1 acceptor vectors are binary vectors and can be used directly for plant transformation to further maximize prototyping speed. MultiGreen is a 1:1 expansion of the original GreenGate architecture’s grammar and has been demonstrated to efficiently assemble plasmids with multiple transcriptional units. MultiGreen has been validated by using a truncated violacein operon from *Chromobacterium violaceum* in bacteria and by deconstructing the RUBY reporter for *in planta* functional validation. MultiGreen currently supports many of our in-house multi transcriptional unit assemblies and will be a valuable strategy for more complex cloning projects.

## Introduction

Simple made-to-order gene expression units met research and commercial requirements for the first few decades of transformation technology. These expression units were assembled with traditional cut-and-paste-style molecular cloning [1]. In principle, the only requirements are a backbone with a multiple cloning site, access to compatible restriction enzymes for the insert, the consumables for PCR and restriction ligation, and the result is a transcriptional unit with a minimum of four components: the promoter and 5’ UTR, coding sequence, terminator and 3’ UTR, and vector backbone [2].

Current needs increasingly require multiplexed guides for editing, or multiple enzymes for metabolic engineering or trait-stacking. While cut-and-paste cloning is efficient and largely the basis for many modern cloning strategies, its utility decreases as the number of transcriptional units to be assembled increases. A more streamlined approach to plasmid construction is the reliance on standardized components and the use of modular assembly.

Gateway cloning is one of the earliest methods for modular assembly and employs site-specific recombination for plasmid construction [3]. Gateway cloning was commercialized by Invitrogen in the 1990s and relies upon recombinase enzymes from phage λ that can recombine DNA sequences flanked by Gateway recombination sites in a series of reactions—the BP and LR reactions recombining between *attB*-*attP* and *attL-attR* sites, respectively. Additional improvements led to MultiSite Gateway cloning, with which two to four DNA fragments can be combined in a single vector using the BP and LR Clonase enzymes [4]. Besides cost, Gateway recombination leaves recombination sites in the final assembly, called scars, that can add 8 to 13 additional amino acids to final reading frames [5].

A more modern approach to modularity is Golden Gate cloning [5]. Golden Gate cloning, which uses one step, one pot, rapid assembly, was proposed initially to address the major limitations of Gateway cloning. It relies on Type IIS restriction enzymes that cleave double-stranded DNA outside of their recognition sequence, leaving behind a minimal overhang. By deploying diverse Type IIS sticky overhangs, multiple DNA fragments can be combined unidirectionally in a one-pot reaction using only one restriction enzyme. As many as 52 parts have been assembled in a one-pot reaction with Golden Gate cloning, with approximately 50% efficiency [6].

There are several different Golden Gate cloning systems available that rely on Type IIS restriction enzymes, i.e., *Aar*I/*Paq*CI, *Eco*31I/*Bsa*I, and *Bsm*BI/*Esp*3I, such as GoldenBraid [7], Mobius Assembly [8], GreenGate [9], and MoClo [10], Each of these systems provides varying leverage over the components necessary for gene expression and approach modularity in a similar fashion, with unique grammar. The grammar of each system is determined by four factors 1) the Type IIS enzymes being used, 2) the coordinated overhangs left behind after Type IIS enzyme digestion for directional assembly, 3) the number of units into which individual transcripts are split into for modularity and 4) organization of the different stages of assembly —such as in parallel, in series, or a combination of the two.

Modern modular assembly systems must fulfill two tasks. The first is assembling promoters, coding sequences, and other components into transcriptional units. Second, the resulting transcriptional units must be assembled into a single multigene vector. Hence, each modular cloning system typically relies on multiple stages of assembly.

The initial stage produces “base,” “entry,” or “level 0” plasmids, each of which contains one component for gene expression, such as a promoter, flanked by Type IIS restriction sites. Subsequent one-pot reactions enable freeing of the components and unidirectional production of individual transcriptional units in another plasmid. Initial assemblies of “level 0” parts are often referred to as “level 1” assemblies and are composed of individual transcriptional units. Level 1 assemblies can be joined together with other level 1 assemblies into a final, “level 2” assembly composed of multiple transcriptional units [7, 8, 10]. Alternatively, level 1 assemblies can be iteratively combined for multiplexing before a final level 2 condensation.

Current modular cloning systems vary in complexity, the exact composition of overhangs, and the Type IIS enzyme responsible for freeing components. Except for GreenGate, all employ multiple Type IIS restriction enzymes to produce a final assembly. While using multiple enzymes can increase the burden of domestication, or the deactivation of naturally occurring Type IIS sites within parts, it enables efficient condensation of multiple transcriptional units either sequentially, or even by “braiding” of assemblies together in a cyclical fashion as in GoldenBraid [7].

GreenGate strikes a balance between ease of use and modularity, enabling control over all the key elements of a cassette for plant expression: promoter, N-tag peptide, coding sequence, C-tag, terminator, and plant selectable marker. Each of these modules are flanked by *Bsa*I sites, leaving seven, unique 4 base pair overhangs for unidirectional assembly—named the A, B, C, D, E, F, and G overhangs. The eighth, optional, H overhang can be introduced through a methylated oligoduplex for multiplexing and is the biggest pitfall of the GreenGate kit.

Therefore, we propose MultiGreen as a multiplexing solution, inspired by the original architecture set forth by GreenGate, to enable intuitive multiplexing while using only the syntax laid forth in the original GreenGate kit. MultiGreen is composed of two approaches to multiplexing: MultiGreen 1.0— assembly in series, and MultiGreen 2.0 assembly in parallel. Multiplexed assembly with MultiGreen is accomplished via a suite of additional vectors that enable condensation of multiple transcriptional units into one final plasmid. Both MultiGreen 1.0 and 2.0 can be combined for more complicated assemblies, or to insert transcriptional blockers [11] between transcriptional units without deviation from the original kit’s grammar. MultiGreen also encompasses a suite of linker modules that bridge gaps in parallel designs that don’t occupy all the conventional GreenGate overhangs, enabling assembly when some level 0 modules are absent. The suite of MultiGreen plasmids will be made available as a kit through Addgene.

## Methods

### General molecular biology reagents and methods

All consumable enzymes for restriction ligation and Gibson assembly were procured from NEB (New England Biolabs, Ipswich, MA) including *Bsa*I-HF-v2 (NEB #R3733S), *Esp*3I (NEB #R0734S), NEBuilder HiFi DNA Assembly Master Mix (NEB #E2621S), NEBridge Ligase Master Mix (NEB # M1100S), and cloning strains of *E. coli*, including DH5α (NEB #C2987H) and 10-beta (NEB # C3019H). For vectors containing the *ccd*B/CmR cassette, One Shot *ccd*B Survival cells (ThermoFisher Scientific, Waltham, MA, #A10460) were used during plasmid propagation.

All plasmids were isolated using standard protocols and reagents from the GenCatch plasmid DNA mini-prep kit (Epoch Life Science, Missouri City, TX). Amplification of target sequences under 5 kb were produced using Q5 high fidelity polymerase (NEB # M0491S) under the recommended three-step amplification conditions, while amplicons over 5 kb were produced using PrimeStar GXL polymerase (Takara Bio, San Jose, CA, #R050A) in a two-step amplification reaction. All NEBuilder HiFi assemblies were executed following standard incubation [12] and transformation procedures [13] into their respective destination strains. All products of amplification were sequenced with either Sanger sequencing (Genewiz, South Plainfield, NJ) for regions <1600 bp, or Oxford Nanopore long-read sequencing (Plasmidsaurus, Eugene, OR) for regions >1600 bp.

### Transformation of bacterial strains

Commercially available cloning strains of *E. coli* were used for transformation of all assemblies. The same protocol was used for all chemically competent strains used, adjusting the resuspension medium, outgrowth duration, and antibiotic selection for the strain and resistances being used. Two µL of assembly mix were combined with 10 µL of freshly thawed competent cells and incubated on ice for 30 mins in a 2-mL Eppendorf tube. The bacteria were then heat shocked at 42°C for 30s and immediately returned to ice for 5 minutes. Two hundred µL of resuspension medium (SOC for DH5α and OneShot cells; NEB 10-beta stable outgrowth medium for 10-beta cells) was added to the tubes.

Plasmids selected on ampicillin were immediately plated on LB medium containing ampicillin while all other antibiotic selections were incubated in a 37°C shaking incubator for a 1-hr outgrowth. *E. coli* transformations were plated on LB supplemented with 15 g·L^-1^ BactoAgar and appropriate antibiotics for the plasmid being selected—100 µg·mL^-1^ ampicillin, 50 µg·mL^-1^ kanamycin, 25 µg·mL^-1^ chloramphenicol, 100 µg·mL^-1^ spectinomycin. Colonies were isolated for liquid culture and sequencing following overnight incubation at 37°C.

### Level 0 entry vector construction

Level 0 entry vectors were created through a scaled down NEBuilder (NEB, Ipswich, MA) reaction. For single insert entry vectors, individual PCR amplicons were produced using Q5 high fidelity polymerase (NEB # M0491S) and isolated from a 0.5X TBE gel by centrifugation of a gel core through an EconoSpin column (Epoch Life Science, Missouri City, TX), at 10000 x *g* for three minutes. One µL of the eluate was immediately combined with 1 µL of *Bsa*I-HFv2-digested level 0 entry vector normalized to 50 ng·µL^-1^ and 1 µL of NEBuilder HiFi DNA Assembly Master Mix. Single-insert assemblies were incubated at 50°C for 20 minutes prior to transformation into chemically competent DH5α cells. A list of primers used for assemblies is in S1 Table.

Domestication of *Esp*3I sites in level 0 inserts was addressed via site-directed mutagenesis with degenerate PCR primers. Briefly, overlapping primers for NEBuilder assembly were designed with a minimum of 15-bp-overlap spanning the restriction site to be domesticated. A single point mutation was introduced within the overlap to disable the site. If the domestication site was within a coding sequence, synonymous mutations were introduced. Following amplification of the two fragments spanning the site, the level 0 entry vector construction proceeded as described with NEBuilder. Components requiring multiple restriction site domestications per insert were incubated at 50°C for 1 hour prior to transformation, rather than 20 minutes. Further, 0.5 µL of NEBuilder HiFi DNA Assembly Master Mix was added for each additional insert.

As the default GreenGate entry vector plasmids contain two *Esp*3I sites within 40 bp of each other in the backbone, we created a derivative suite of base level 0 entry vectors eliminating the *Esp*3I sites. Domestication of these sites is not essential for successful MultiGreen assembly using pre-made level 0 entry vectors, as the overhangs resulting from *Esp*3I digestion of the entry vector backbone are incompatible with the overhangs for GreenGate assembly. Initial attempts of single-stranded oligo assembly of the digested vector failed, so we combined inverse PCR with single-stranded oligo assembly to disable the two *Esp*3I sites. The sequences 5’-AGACGAAAGGGCCTCGTGAT-3’ and 5’-AGACGGTCACAGCTTGTCTG-3’ were used as the forward and reverse primers, respectively, to amplify the entire entry vector using Q5 high fidelity polymerase. The amplicon was isolated as described for level 0 amplicons and combined with 1 µL of a 1:250 dilution of 100 µM single-stranded oligo for recircularization (5’-CAGACAAGCTGTGACCGTCTCCGGGAGCTGCATGTGTCAGAGGTTTTCACCGTCATCACCGAAACGCGCGAGACGAAAGGGCCTCGTGAT-3’). One µL of NEBuilder HiFi DNA Assembly Master Mix was then added, and the mix incubated for 1 hour at 50°C before transformation into *E. coli* strain DB3.1.

Level 0 entry clones for linker modules not requiring PCR were assembled via a sense/antisense single stranded oligo NEBuilder hybridization. Briefly, single-stranded oligos were designed to contain 20 bp of complementarity to their desired vector backbone, the required Type IIS recognition sequence and overhang, and 15-20 bp of complementarity to the 5’ oligo for hybridization. Then, 1:250 dilutions of 100 µM oligos (Millipore Sigma, Burlington, MA) were prepared with type I water. One µL each of the sense and antisense oligo dilutions were then combined with 1 µL of cut level 0 entry vector at 50 ng·µL^-1^ and 2 µL of NEBuilder HiFi DNA Assembly Master Mix. The reaction mix was then incubated at 50°C for 1 hour prior to transformation. A list of oligos used for producing linker modules can be found in S1 Table.

### Level 1 MultiGreen intermediary vector construction

Level 1 MultiGreen intermediary vectors were made through successive PCR amplification and assembly using NEBuilder. Destination vector pGGP-AG [14] was domesticated of its three *Esp*3I sites as described, generating pVP076, which was then used as the template in a second round of PCR amplification and NEBuilder assemblies to generate the initial suite of MultiGreen level 1 acceptor vectors.

Six sets of overlapping PCR primers were designed to include both homology to the backbone, pairs of flanking *Esp*3I and *Bsa*I sites, GreenGate-compatible overhangs, and homology to the *ccd*B/chloramphenicol cassette to retain the counterselection used in all GreenGate base vectors. Plasmid VP076 was digested in a 10 µL digest using 1 µg of plasmid, 0.5 µL of *Bsa*I-HF-v2, and 1 µL of CutSmart Buffer (NEB #B6004S) for one hour at 37°C. Following incubation, 2 µL of loading dye were added, the digestion was run on a gel, and the backbone isolated as before. One µL each of PCR amplicon, backbone, and NEBuilder were assembled and transformed into OneShot *ccd*B survival cells as outlined above, generating pVP078, pVP079, pVP080, pVP081, pVP082, and pVP083 corresponding to MultiGreen level 1 acceptor vectors with final GreenGate-compatible AB, BC, CD, DE, EF, and FG overhangs, respectively.

To make another destination vector, and progenitor to a second suite of MultiGreen intermediary vectors, kanamycin resistance was introduced using restriction ligation of pGGP-AG and NEBuilder assembly with an amplified aph(3′)-Ia kanamycin resistance gene from p201NCas9 [15]. Destination vector pGGP-AG was digested as above, except with *Ava* II instead of *Bsa*I, ejecting a 641-bp fragment of the spectinomycin resistance gene from the vector. The kanamycin resistance gene was then amplified using 5’-CGTATGCGCTCACGCAACTGGATGAGCCATATTCAACGG-3’ as the forward primer and 5’-AAAGAGTTCCTCCGCCGCTGTTAGAAAAACTCATCGAGC-3’ as the reverse primer, sharing a 20-bp overlap with the cut vector, and maintaining the reading frame established by the remaining 5’ end of the spectinomycin resistance gene. The NEBuilder assembly was performed and transformed into OneShot *ccd*B survival cells, and selected on LB kanamycin for successful assembly, generating pGGPK-AG2.

The *Esp*3I sites of pGGPK-AG2 were domesticated as in pVP076, with an additional pair of overlapping degenerate primers for the *Esp*3I site within the kanamycin resistance gene to generate pVP077. Plasmid VP077 was subsequently used for its kanamycin-resistant and *Esp*3I-domesticated backbone to generate a second suite of MultiGreen level 1 acceptor vectors, pVP311, pVP312, pVP313, pVP314, pVP315, pVP316, following the same steps and amplifications as with pPV078-pVP083. pVP076 was also digested and assembled via NEBuilder with a PCR amplicon containing mRFP1E chromoprotein (Addgene #160442) in place of *ccd*B/CmR, generating pVP096.

### Level 1 and Level 2 GreenGate assembly with NEBridge

Level 1 and level 2 assemblies were performed using NEBridge ligase master mix described previously unless otherwise noted [16]. Level 1 assemblies with solely GreenGate level 0 entry vectors require only *Bsa*I as the restriction enzyme. Assemblies using previously made MultiGreen level 1 assemblies also required *Esp*3I in the reaction cocktail to facilitate freeing of level 1 modules. Recurrent level 1 and level 2 assemblies were performed in a one-pot reaction with a total volume of 15 µL. Each overhang in the assembly was be used, either directly through the compatible level 0 module or level 1 assembly, or indirectly via a MultiGreen linker module spanning the unused overhangs.

### MultiGreen cloning validation through protoviolaceinate biosynthesis

To ensure MultiGreen functioned as envisioned, protoviolaceinate biosynthesis from *Chromobacterium violaceum* was reconstructed as a four transcriptional unit stack using MultiGreen 2.0. Level 0 components were produced as outlined above and as listed in S2 Table. Initial attempts at level 1 assemblies sought to combine each violacein gene with the strongest in house characterized promoter pGG-A-PJ23119:PGLPT-C/pVP217, pGG-D-rrnbT1-F/pVP223 terminator into spectinomycin resistant MultiGreen 2.0 intermediary vectors AB/pVP078, BC/pVP079, DE/pVP081, and EF/pVP082, respectively. In addition, eGFP was assembled with the strongest characterized promoter into a spectinomycin resistant MultiGreen 2.0 intermediary FG/pVP083.

Plasmid MG2.0-B-PJ23119:PGLPT:vioB:rrnb1-C/pVP268, pMG2.0-E-PJ23119:PGLPT:vioE:rrnb1-F/pVP271, and pMG2.0-F-PJ23119:PGLPT:gfp:rrnb1-G/pVP272 were successfully cloned. Plasmid MG2.0-A-PJ23119:PGLPT:vioA:rrnb1-B/pVP267 and pMG2.0-D-PJ23119:PGLPT:vioD:rrnb1-E/pVP270, however, failed to be isolated after repeated assembly attempts.

Given the failed assembly of *vioA* and *vioD*, weaker promoter variants of each *vioA—*pMG2.0-A-PJ23100:B0030:vioA:rrnb1-B/pVP250 , and *vioD*—pMG2.0-D-PJ23100:B0030:vioD:rrnb1-E/pVP253 were cloned. The weakly expressing *vioA* and *vioD* were then combined with the strongly expressing *vioB* and *vioC*, pGG-bd-C-dummy-D /pVP293, pMG2.0-F-PJ23119:PGLPT:gfp:rrnb1-G/pVP272 , and kanamycin-resistant pVP096 in a level 2 MultiGreen assembly to produce pVP290— expressing all four genes necessary for protoviolaceinate biosynthesis in *E. coli*. pVP250 is a very low-yielding plasmid (50-60 ng·µL^-1^). Therefore, the NEBridge assembly was performed as outlined above using only 0.02 pmol of each input component.

Three biological replicates of the MultiGreen 2.0 assemblies were constructed and transformed on independent days into NEB 10-beta chemically competent cells. Three serial dilutions of the bacterial culture were plated on LB kanamycin plates and incubated for 40 hours prior to imaging to allow for pigment production. Serial dilutions with discernable colony separation were used for imaging and assembly efficiency quantification. Undigested destination vector carryover (pVP096) was scored for fluorescence of the mRFP1E visual reporter situated between assembly sites. Three random colonies failing to express both the protoviolaceinate pigment and the mRFP1e reporter were sent for whole plasmid sequencing via Plasmidsaurus to evaluate assembly fidelity. Colony counts were tallied, and efficiency calculated in Microsoft Excel for Mac version 16.81 (Microsoft Corporation, Redmond, WA).

### Deconstructed RUBY Nicotiana benthamiana assay

To further test MultiGreen in a eukaryotic system, the enzymes of the RUBY reporter [17] were reassembled using separate MultiGreen 2.0 level 1 assemblies for each transcriptional unit. Level 1 assemblies were performed with NEBridge Ligase Master Mix using the following level 0 components: pGG-A-GmUbi3L-B containing the *GmUbi3L* promoter from *Glycine max* [18], pGGB003 [9], pGG-C-CYP76AD1-D/pVP214 and pGG-C-DODA-D/pVP215 and pGG-C-GT-D/pVP216 modules corresponding to each coding sequence in the RUBY reporter [17], pGGD002 [9], pGG-E-StPinII-F containing the *PinII* terminator from *Solanum tuberosum* [19], pGG-F-dummy-G. CYP76AD1, DODA, and GT were assembled into the spectinomycin-resistant MultiGreen2.0 AB/pVP078, BC/pVP079, and CD/pVP080 intermediaries, respectively. Correct clones were then used in a level 2 MultiGreen assembly using NEBridge Ligase master mix with pGG-C-dummy-D/pVP121 linker into the kanamycin-resistant pVP096 domesticated destination vector.

Competent cells of an in-house disarmed *thyA*-*recA*-variant of *Agrobacterium rhizogenes* strain K599 (aka NCPPB2659)[20] were prepared as specified in the BioRad MicroPulser™ (Biorad Laboratories, Hercules CA, US) manual. Cells were recovered in liquid YP medium [21] containing 150 mg·L^-1^ thymidine and no antibiotics for 4 h at 28°C with shaking. After recovery, the bacterial suspension was plated onto solid YP medium containing 150 mg·L^-1^ thymidine and 50 mg·L^-1^ kanamycin. Single kanamycin-resistant colonies growing on the electroporation plate were inoculated into 25 mL liquid YP cultures for overnight incubation at 28°C with shaking. *Agrobacterium* suspensions and *Nicotiana benthamiana* infiltrations were conducted as described previously [11], only deviating by addition of thymidine to all media containing *Agrobacterium*. The three youngest, expanded leaves were infiltrated with bacterial suspension of OD600 of 0, 0.1, 0.2, 0.5, and 1.0.

Three days after infiltration, betanin production was quantified based on previously outlined experimental procedures [22]. Individual 8-mm leaf discs were collected using disposable biopsy punches, one leaf per infiltration site, three leaves per plant, and added to 2-mL Eppendorf tubes containing 2 mL of 100% ethanol for chlorophyll bleaching for a minimum of 24 h. Following bleaching, leaf discs were individually removed, rinsed with fresh 100% ethanol, and added to a 96-well plate (VWR, Radnor, PA, # 10861-562) containing 150 µL of type I water per well. The 96-well plates were allowed to incubate on the bench top for 24 hours, upon which leaf discs were removed and absorbance at 538 nm was measured on a Biotek Synergy2 plate reader (BioTek Instruments Inc., Winooski, VT, USA). Absorbance readings were imported to GraphPad Prism 10 for macOS version 10.2.0 for plotting and analysis.

## Results and discussion

### MultiGreen 1.0 – Multiplexing in series

The original GreenGate cloning kit is composed of a suite of entry modules containing convergent pairs of *Bsa*I sites flanking components of interest for one-pot assembly [9]. For Golden Gate assembly reactions, it is important to domesticate, i.e., remove all the unnecessary restriction sites for the Type IIS enzyme used for assembly to avoid unintended mis-assemblies. Fortunately, many of the Type IIS restriction enzymes have recognition sequences >5 bp, reducing their frequency of occurrence.

Often, fragments for Golden Gate assemblies are plasmids containing Type IIS sites flanking the insert, or PCR amplicons with compatible cut sites within or near the amplification primers. If plasmids are used as the DNA source, it is common to use alternating antibiotic selections between the parent and child vectors to circumvent carryover, or persistence, of the parent plasmid in subsequent reactions [7, 9, 10, 23].

The canonical entry vectors for GreenGate cloning include A, B, C, D, E, F, and G overhangs in ampicillin-resistant level 0 plasmids to enable unidirectional assembly (Fig 1 A). Multiplexing as proposed with the original GreenGate kit introduces the optional H overhang via methylated oligos and linker modules between F and G, thereby multiplexing in series from a 5’→ 3’ direction. For duplex assemblies with GreenGate, one additional overhang can be introduced with a specifically methylated oligoduplex, such that the first transcriptional unit assembled becomes the destination vector for the second transcriptional unit by exploiting the methylation sensitivity of *Bsa*I, effectively blocking digestion of the hidden overhangs in the first iteration of cloning [9]. Multiplexing via the methylated oligoduplex strategy is not only cost-prohibitive, but results in a heightened screening burden, as there is no intrinsic counterselection between assemblies.

**Fig 1.**
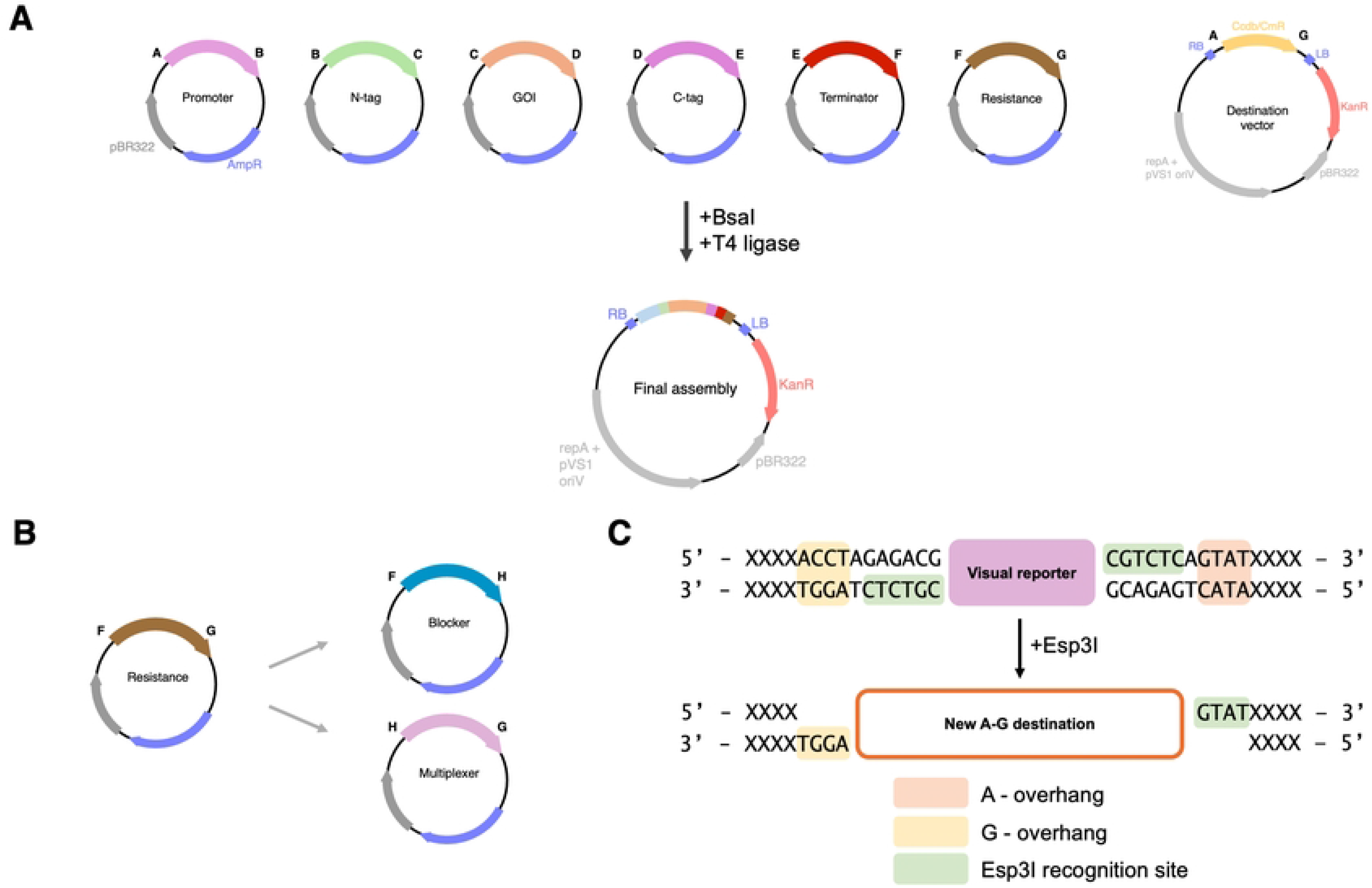
MultiGreen 1.0 - multiplexing in series. A) Schematic overview of a standard GreenGate reaction condensing six entry vectors, AB Promoter, BC N-tag, CD GOI, DE C-tag, EF terminator, FG resistance, into a destination vector using a one-pot *Bsa*I mediated restriction ligation reaction. B) MultiGreen 1.0 expansion modules introduce the H overhang as a split of two new Level 0 entry vectors, the FH module is reserved for transcription blockers and the HG module is reserved for the multiplexer. The HG multiplexer module contains a pair of *Esp*3I sites internal to the *Bsa*I sites for conventional GreenGate assembly, but external to the chromoprotein reporter. Each assembly that incorporates the level 0 HG and FH modules for multiplexing can be visually selected for integration of the multiplexer module. Subsequent MultiGreen 1.0 reactions should alternate visual reporters to efficiently screen one assembly round from the next as the chemical selection is set by the initial destination vector. The assembly for the final transcriptional unit should either include the FH transcription blocker and HG filler modules to terminate the multiplexing, or alternatively a level 0 FG plant selectable marker module can be included. C) Detailed view of how the multiplexer operates by incorporating an external set of *Esp*3I sites flanking a visual reporter. Inclusion of the FH and HG modules *in lieu* of the FG followed by selection for colonies expressing the visual reporter enables iterative stacking in series.

As an alternative, MultiGreen 1.0 introduces the additional H overhang via two new Level 0 entry vectors, the FH and HG modules (Fig 1 B). This approach is possible by incorporating a second Type IIS restriction enzyme, *Esp*3I. Therefore, all components in an assembly must be domesticated for *Esp*3I or adequately screened for correct assembly from one reaction to the next. Domestication of existing components can be readily performed by inverse PCR with degenerate primers destroying the undesired cut site followed by restriction-ligation or Gibson assembly. Alternatively, single-stranded oligos have been successfully used in conjunction with digested vectors in a Gibson assembly reaction for site domestication in an oligo-stitching reaction [24].

Like *Bsa*I, *Esp*3I is has a 6-base pair recognition sequence, leaving a 4-base pair sticky overhang. The MultiGreen 1.0 FH module is reserved for a transcription blocker, or alternatively can contain a spacer sequence (Fig 1 B). We preconfigured several HG multiplexer modules with visually distinct chromoprotein reporters including meffBlue [25], mRFP1e [26], and spisPink [25]. With each iteration of multiplexing, correctly assembled clones possessing the multiplexer module for the next round of assembly can be selected for expression of the respective chromoprotein reporter. The availably of multiple reporters enables recurrent passage of assemblies through visual screening for the reporter included in each assembly step. The chromoproteins that come with MultiGreen 1.0 can be swapped for other reporters, or even antibiotic resistance genes by performing a standard PCR and either restriction-ligation or Gibson assembly.

The chromoproteins within the HG multiplexer were amplified directly from Addgene bacterial expression plasmids. The chromoproteins in the HG multiplexer modules are flanked by convergent *Bsa*I sites, enabling assembly alongside other level 0 entry vectors, as is the case in a conventional GreenGate reaction. Just internal to the *Bsa*I sites, but still external to the chromoprotein, are a pair of divergent *Esp*3I sites. These remain undigested in the initial assembly, as *Bsa*I is the only restriction enzyme present, effectively providing a future pair of compatible A-G destination overhangs in a subsequent reaction with *Esp*3I (Fig 1 C). A correct clone expressing the chromoprotein can then be used as the destination vector for a subsequent assembly by digestion with *Esp*3I, freeing the receptive destination A-G overhangs for another transcriptional unit assembly.

MultiGreen 1.0’s time for assembly scales at an approximate rate of 2n +1 days, where n is the number of transcriptional units, and assumes all level 0 components and reagents are available when assembly is started, and that diagnostic screens occur the same day colonies appear. This rate of assembly is tolerable for simple assemblies but draws out when additional transcriptional units are desired. A unique advantage of MultiGreen 1.0 is that if there is a gene cassette containing *Esp*3I sites that cannot be easily domesticated, such as in regulatory sequences that may need additional validation after domestication, assembly can be structured such that the non-*Esp*3I domesticated components are in the final reaction, where only *Bsa*I is the active restriction enzyme.

### MultiGreen 2.0 – Multiplexing in parallel

While MultiGreen 1.0 enabled efficient transcriptional unit stacking following the GreenGate multiplexing grammar, its efficiency of assembly lags for constructs with many transcriptional units. For example, assembling six cassettes with MultiGreen 1.0 would require a minimum of 13 days of cloning to produce the one final vector. Additionally, since MultiGreen 1.0 is multiplexing in series, and each successive reaction becomes the destination vector of the next, if it is desirable to exchange an individual transcriptional unit after assembly, it would be necessary to revert to the assembly just before the unwanted unit and restart incorporating the remaining transcriptional unit. To circumvent the limitations of overall time efficiency of assembly, maximize the compatibility of level 1 assemblies across constructs, and maintain compatibility with the original GreenGate syntax, we propose MultiGreen 2.0 – multiplexing in parallel (Fig 2).

**Fig 2.**
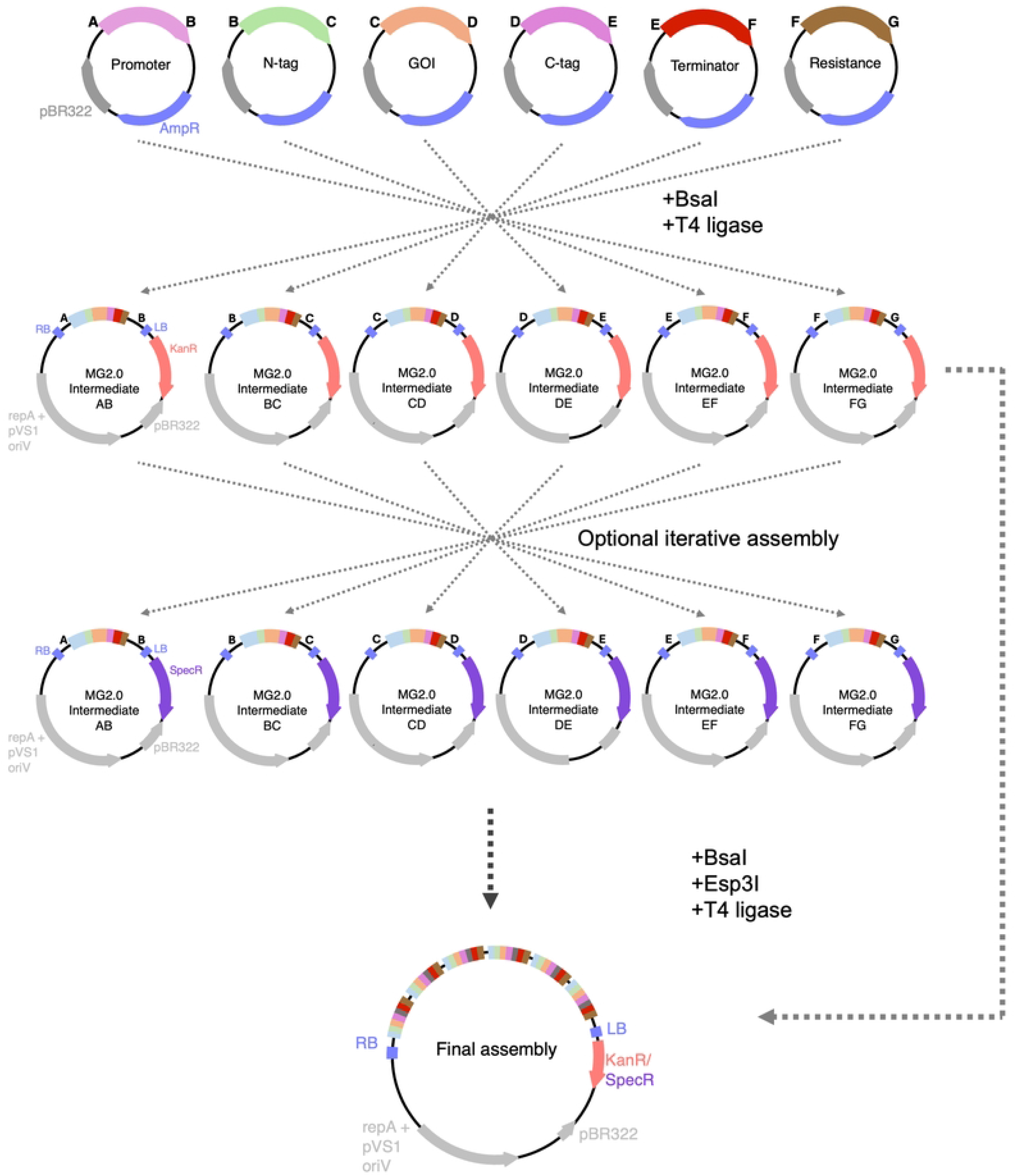
MultiGreen 2.0 - multiplexing in parallel. A schematic overview of multiplexing in parallel with MultiGreen 2.0. MultiGreen 2.0 uses a suite of level 1 acceptor vectors to produce initial concatenations of entry vectors. The level 1 acceptor vectors are not ampicillin resistant, allowing for chemical counterselection from level 0 entry vectors. Level 1 acceptor vectors are binary vectors themselves and can be used directly in *Agrobacterium* for (co)infection. Up to 6 Level 1 assemblies can be combined per reaction. By alternating selection choice of level 1 acceptor vectors, assemblies can be iteratively combined in multiples of 6. Note that alternating selection of level 1 acceptor vectors is only necessary for assemblies with >6 transcriptional units. Conventional GreenGate level 0 entry modules can be incorporated in any level 2 assembly with compatible overhangs, provided both Type IIS enzymes are included in the one-pot reaction cocktail.

MultiGreen 2.0 takes inspiration from the other modular cloning standards that support multiplexing, such as GoldenBraid [7], Mobius Assembly [8], and MoClo [10] by parallelizing multiplexing with a suite of intermediary level 1 acceptor vectors. MultiGreen 2.0 assemblies are performed into a suite of level 1 acceptor vectors mirroring the overhang syntax of GreenGate assembly, allowing for concatenation of up to six level 1 assemblies in parallel. Alternation of antibiotic selection allows this to occur indefinitely with adequate planning of which level 1 acceptor vector complement to use, and strategic reversion to level 0 GreenGate entry vectors. Using the same assumptions as in MultiGreen 1.0 for assembly constraints, MultiGreen 2.0 assembly time scales at a rate of 2*⌈*log*_6_*n*⌉+3 days, where *n* is the number of transcriptional units. When comparing the time efficiency between MultiGreen 1.0 and 2.0, that is a time savings of nearly two weeks for a six transcriptional unit assembly, taking a minimum of 5 days with MultiGreen 2.0. By cloning with MultiGreen 2.0, the assembly is also better suited for future proofing, since transcriptional units can be more easily exchanged later.

The level 1 intermediary vectors of MultiGreen 2.0 use an additional pair of flanking *Esp*3I sites to free assembled transcriptional units for additional rounds of assembly (Fig 3). Each MultiGreen 2.0 level 1 acceptor vector contains a pair of *Esp*3I sites externally flanking the *Bsa*I sites necessary for a GreenGate single transcriptional unit assembly. When a GreenGate reaction is performed using a MultiGreen 2.0 level 1 acceptor vector as the destination vector, it can then be digested with *Esp*3I to liberate the assembled transcriptional unit for another GreenGate reaction, since *Esp*3I digestion of MultiGreen 2.0 level 1 assemblies leaves behind either AB, BC, CD, DE, EF, or FG overhangs, depending on the MultiGreen level 1 intermediary destination vector chosen (Fig 2). The freed overhangs enable level 1 assembly products to act as pseudo-level 0 modules.

**Fig 3.**
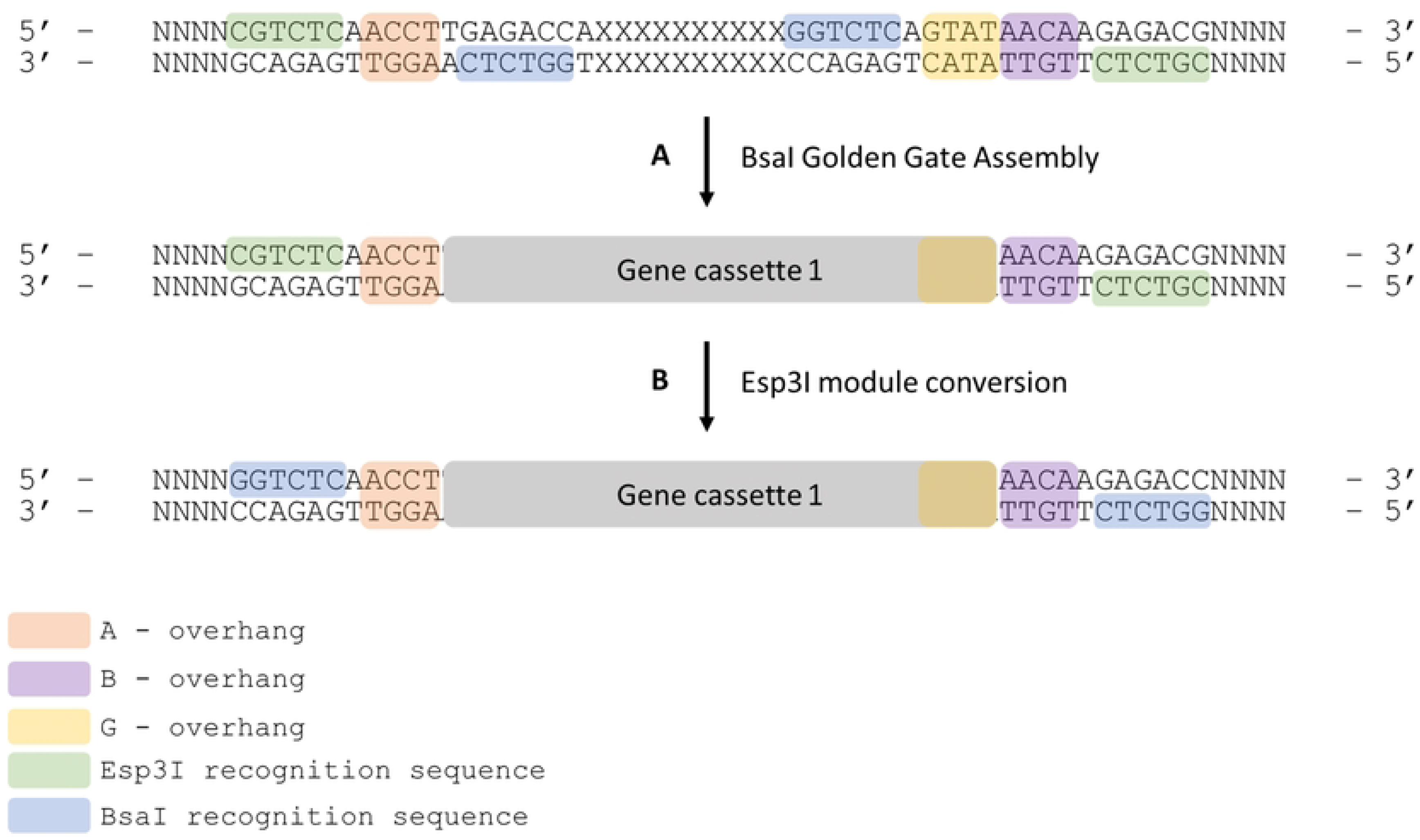
Precise mechanism of MultiGreen 2.0 cloning and reversion to level 0 modules from level 1 assemblies. A series of level 1 acceptor vectors are made with complimentary overhangs for each GreenGate entry module (AB, BC, CD, DE, EF, FG). A) The above example is for a MG2.0 AB assembly. A standard GreenGate cloning reaction is performed using the MG2.0 AB intermediary vector as denoted in A. The product of that assembly will hold the first gene cassette in the series. By the nature of the flanking Type IIS restriction enzyme sites, the recognition sequence for *Bsa*I is dropped out, but the 4bp overhangs are retained. This intermediary MG2.0 AB vector containing Gene cassette 1 can be used directly in a level 2 assembly incorporating Esp3I, be digested and used as an AB fragment in a subsequent GreenGate reaction or may be B) ligated into a GreenGate AB entry vector backbone effectively converting the level 1 assembly into a level 0 module.

At its core, MultiGreen 2.0 enables recursive assembly of GreenGate modules. In other words, it enables performing GreenGate reactions into MultiGreen level 1 acceptor vectors, which when digested with *Esp*3I, function as level 0 parts (Fig 2). Should transcription blockers be a desirable feature between stacked gene cassettes, the dedicated FH transcription blocker module and HG filler module from MultiGreen 1.0 can be incorporated in each level 1 assembly, resulting in transcription blockers or other spacers being placed between cassettes. The HG filler module allows for GreenGate assembly with the additional H overhang when multiplexing in parallel.

In MultiGreen version 2.0, level 1 acceptor vectors are derived from the pVS1 backbone of pGGP-AG [14] for use as binary vectors directly in *Agrobacterium*, and each complement of level 1 intermediary vectors has a different antibiotic marker—either spectinomycin or kanamycin. Two antibiotic complements enable the cycling of level 1 assemblies. By alternating kanamycin and spectinomycin selection between reactions, undigested plasmid carryover from one assembly to the next is eliminated. Since *Esp*3I digestion of level 1 MultiGreen 2.0 assemblies frees level 0-compatible overhangs, the individual transcriptional units released during *Esp*3I digestion can also be readily converted back to true level 0 modules with T4 ligase. Should a level 0 reversion be desired for a particular assembly, it can be efficiently performed as outlined in S1 Method. A list of all plasmids in the MultiGreen kit can be found in Table 1.

**Table 1.** MultiGreen Kit Components. Includes all vectors for MultiGreen 1.0 cloning In series, MultiGreen 2.0 cloning in parallel, and adapters.

| Shorthand | Verbose | Selection | Description |
| --- | --- | --- | --- |
| pVP076 | pGGP-AG-e | Spec | pGGP-AG fully domesticated of *Esp3I* sites |
| pVP077 | pGGPK-AG2-e | Kan | pGGPK-AG2 fully domesticated of *Esp3I* sites |
| pVP078 | pMG2.0-AB-s | Spec | MG2.0 AB level 1 acceptor vector, spectinomycin selection |
| pVP079 | pMG2.0-BC-s | Spec | MG2.0 BC level 1 acceptor vector, spectinomycin selection |
| pVP080 | pMG2.0-CD-s | Spec | MG2.0 CD level 1 acceptor vector, spectinomycin selection |
| pVP081 | pMG2.0-DE-s | Spec | MG2.0 DE level 1 acceptor vector, spectinomycin selection |
| pVP082 | pMG2.0-EF-s | Spec | MG2.0 EF level 1 acceptor vector, spectinomycin selection |
| pVP083 | pMG2.0-FG-s | Spec | MG2.0 FG level 1 acceptor vector, spectinomycin selection |
| pVP084 | pmeffBlue HG | Amp | meffBlue chromoprotein in a MG1.0 HG entry vector |
| pVP085 | pSpisPink HG | Amp | spisPink chromoprotein in a MG1.0 HG entry vector |
| pVP086 | pmRFP1e HG | Amp | mRFP1e chromoprotein in a MG1.0 HG entry vector |
| pVP088 | pGG-H-dummy-G | Amp | MG1.0 HG entry vector containing filler sequence |
| pVP089 | pGG-H-ccdb/CmR-G | Amp | MG1.0 base HG entry vector |
| pVP090 | pGG-F-dummy-H | Amp | MG1.0 FH entry vector containing filler sequence |
| pVP091 | pGG-F-ccdb/CmR-H | Amp | MG1.0 base FH entry vector |
| pVP096 | pGGPK-AG2-mRFP1e-bbe | Kan | pGGPK-AG2 where the ccdb/CmR has been swapped for mRFP1e; Esp3I domesticated backbone |
| pVP333 | pGG-A-dummy-B | Amp | GreenGate AB bridge module containing filler sequence |
| pVP334 | pGG-F-dummy-G | Amp | GreenGate FG bridge module containing filler sequence |
| pVP120 | pGG-B-dummy-G | Amp | GreenGate BG bridge module containing filler sequence |
| pVP121 | pGG-C-dummy-G | Amp | GreenGate CG bridge module containing filler sequence |
| pVP122 | pGG-D-dummy-G | Amp | GreenGate DG bridge module containing filler sequence |
| pVP123 | pGG-E-dummy-G | Amp | GreenGate EG bridge module containing filler sequence |
| pVP183 | pGG-D-dummy-F | Amp | GreenGate DF bridge module containing filler sequence |
| pVP184 | pGG-E-dummy-F | Amp | GreenGate EF bridge module containing filler sequence |
| pVP185 | pGG-B-dummy-F | Amp | GreenGate BF bridge module containing filler sequence |
| pVP186 | pGG-C-dummy-F | Amp | GreenGate CF bridge module containing filler sequence |
| pVP224 | pGG-A-dummy-C | Amp | GreenGate AC bridge module containing filler sequence |
| pVP225 | pGG-A-dummy-D | Amp | GreenGate AD bridge module containing filler sequence |
| pVP226 | pGG-A-dummy-E | Amp | GreenGate AE bridge module containing filler sequence |
| pVP227 | pGG-A-dummy-F | Amp | GreenGate AF bridge module containing filler sequence |
| pVP228 | pGG-B-dummy-D | Amp | GreenGate BD bridge module containing filler sequence |
| pVP229 | pGG-B-dummy-E | Amp | GreenGate BE bridge module containing filler sequence |
| pVP230 | pGG-C-dummy-D | Amp | GreenGate CD bridge module containing filler sequence |
| pVP231 | pGG-C-dummy-E | Amp | GreenGate CE bridge module containing filler sequence |
| pVP293 | pGG-bd-C-dummy-D | Amp | GreenGate CD bridge module containing filler sequence, backbone domesticated of Esp3I sites |
| pVP294 | pGG-bd-C-dummy-D | Amp | GreenGate DE bridge module containing filler sequence, backbone domesticated of Esp3I sites |
| pVP295 | pGG-bd-A-ccdb/CmR-B | Amp | GreenGate AB base module with a Esp3I domesticated backbone |
| pVP296 | pGG-bd-B-ccdb/CmR-C | Amp | GreenGate BC base module with a Esp3I domesticated backbone |
| pVP297 | pGG-bd-C-ccdb/CmR-D | Amp | GreenGate CD base module with a Esp3I domesticated backbone |
| pVP298 | pGG-bd-D-ccdb/CmR-E | Amp | GreenGate DE base module with a Esp3I domesticated backbone |
| pVP299 | pGG-bd-E-ccdb/CmR-F | Amp | GreenGate EF base module with a Esp3I domesticated backbone |
| pVP300 | pGG-bd-F-ccdb/CmR-G | Amp | GreenGate FG base module with a Esp3I domesticated backbone |
| pVP311 | pMG2.0-AB-k | Kan | MG2.0 AB level 1 acceptor vector, kanamycin selection |
| pVP312 | pMG2.0-BC-k | Kan | MG2.0 BC level 1 acceptor vector, kanamycin selection |
| pVP313 | pMG2.0-CD-k | Kan | MG2.0 CD level 1 acceptor vector, kanamycin selection |
| pVP314 | pMG2.0-DE-k | Kan | MG2.0 DE level 1 acceptor vector, kanamycin selection |
| pVP315 | pMG2.0-EF-k | Kan | MG2.0 EF level 1 acceptor vector, kanamycin selection |
| pVP316 | pMG2.0-FG-k | Kan | MG2.0 FG level 1 acceptor vector, kanamycin selection |

### MultiGreen 2.0 cloning validation in bacteria

As a proof of concept for MultiGreen 2.0 and to quantify the efficiency of a dual restriction-enzyme assembly, we reconstructed the partial violacein operon from *Chromobacterium violaceum* for production of protoviolaceinate in *E. coli. C. violaceum* is a species of gram-negative bacterium present various tropical and subtropical areas of the world [27]. A unique feature of the bacterium is that it produces the purple antimicrobial compound, violacein. Violacein is the end-product of a five-enzyme cascade beginning with L-tryptophan. Truncations of this operon result in visually distinct pigments [28]. Previously, Mobius assembly used protoviolaceinate as a functional validation of multiplexing, but only in validation of a level 1 reassembly of the polycistron [28].

The partial violacein operon from *Chromobacterium violaceum* was reassembled to produce protoviolaceinate via four independent transcriptional units using MultiGreen 2.0 (Fig 4 B). To simplify the level 1 assemblies, level 0 modules were constructed with AC, CD, DF, FG overhangs representing the promoter, coding sequence, terminator, and filler sequence. Simplification of the modules was performed to avoid interruption of the gene products or distances between ribosome binding site, transcriptional start site, and start codon for each of the transcripts. Further, initial challenges with producing level 1 assemblies for *vioA* and *vioD* motivated the characterization of four promoter variants using a GFP expression assay (S1 Fig). Of the four promoter variants characterized, one (pGG-A-PJ23119::PGLPT-C/pVP217) highly expresses eGFP relative to that of a DH5α negative control, while the remaining three all express eGFP at low levels relative to that of the DH5α control (S1 Fig).

**Fig 4.**
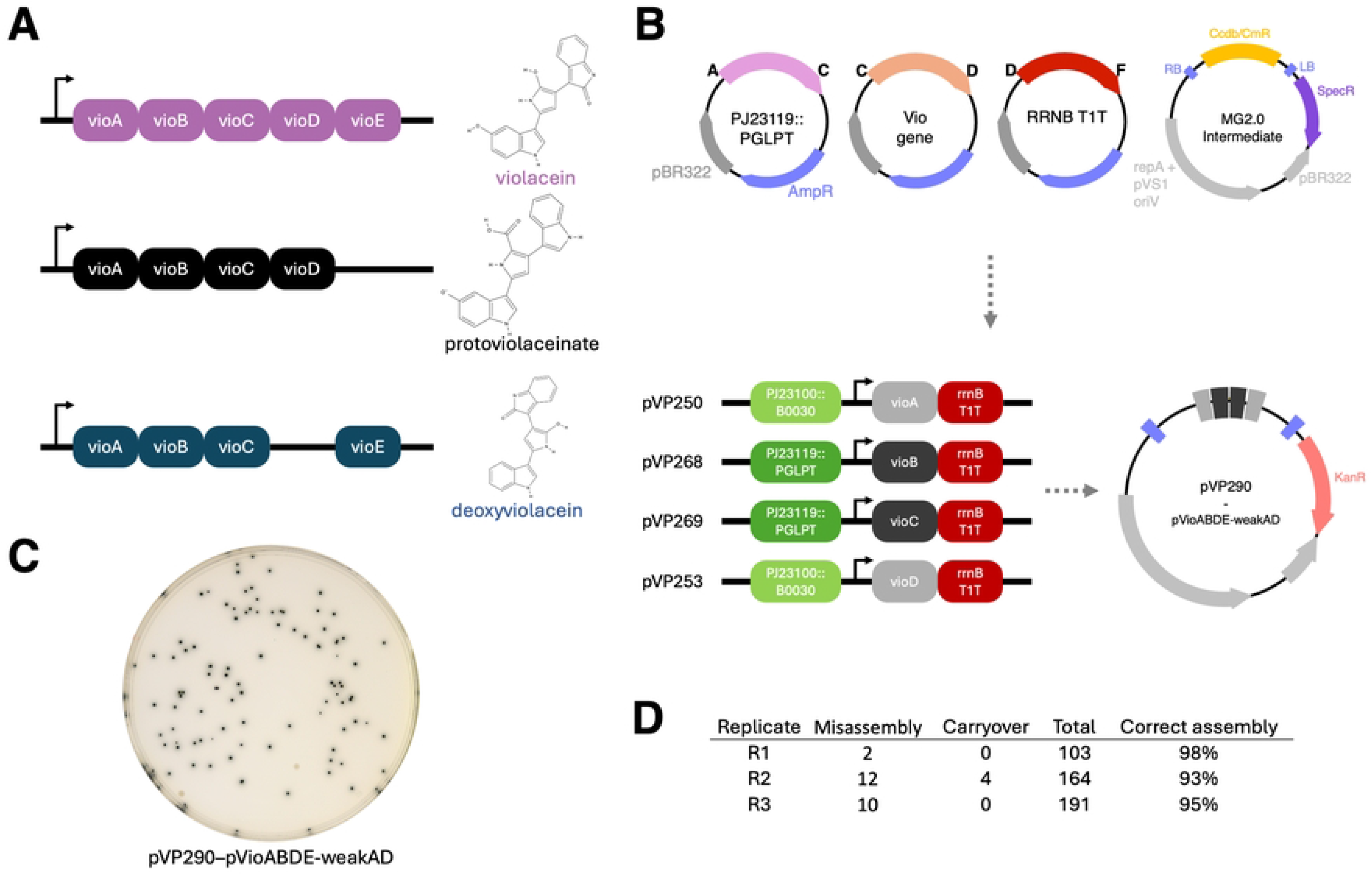
MultiGreen 2.0 cloning validation through protoviolaceinate biosynthesis. A) The full violacein operon and its product alongside two truncations producing different pigmented metabolites. B) Example Level 0 entry vectors for assembling the vioABDE operon using MultiGreen 2.0. Not pictured are the filler sequences to bridge overhang gaps (FG filler for level 1; FG eGFP cassette and CD filler for Level 2). Two different synthetic promoters and the bacteriophage T1 *rrnB* terminator were chosen to drive expression of the four genes from *Chromobacterium violaceum* to produce protoviolaceinate. C) A representative DH10B plate of the Level 2 assembly producing protoviolaceinate as noted by the dark colonies. D) Replicated colony counts of the assembly in independent one-pot reactions produced on different days.

Overall efficiency of the parallelized MultiGreen 2.0 assembly for production of protoviolaceinate using both *Esp*3I and *Bsa*I to concatenate level 1 modules was 95 ± 3% across three replicates (Fig 1 D) as noted by the percentage of colonies expressing the pigment relative to the total of non-carryover colonies. Misassemblies that occurred appeared to consist of escapes resulting in vectors that lack the full *vioABDE* complement (S3 Table), potentially indicating a metabolic cost of *vioABDE* expression in *E. coli* in this fashion. Further, correctly assembled colonies required approximately 40 h for growth to reach a size comparable to that of an overnight *E. coli* colony without the *vioABDE* assembly.

### In planta validation through deconstruction of the RUBY reporter

To further validate MultiGreen as a viable strategy to stack genes in one construct for plant transformation, we deconstructed the RUBY visual reporter [17] using MultiGreen 2.0. RUBY is a three-gene polycistronic transcript, encoding three separate peptides on one transcript. RUBY reconstitutes the biosynthetic pathway from *Beta vulgaris* that converts L-tyrosine into betanin. Each enzyme in the polycistron is separated by self-cleaving P2A peptides [17], using only one promoter and terminator to coordinate expression of all three genes simultaneously. Through a series of MultiGreen parallelized assemblies, we deconstructed the polycistronic transcript for independent transcript expression.

The individual transcriptional units of RUBY were combined with the *Gm*Ubi3L promoter and the StPinII terminator into spectinomycin-resistant MultiGreen 2.0 level 1 acceptor vectors. They were later concatenated with a linker module to produce the final reconstituted RUBY marker in a kanamycin-resistant pVS1-replicon destination vector (Fig 5 A) and evaluated in *Nicotiana benthamiana* leaf infiltration (Fig 5 C). The results of these infiltrations support MultiGreen 2.0 being an effective strategy to assemble multiplexed plasmids for *in planta* expression.

**Fig 5.**
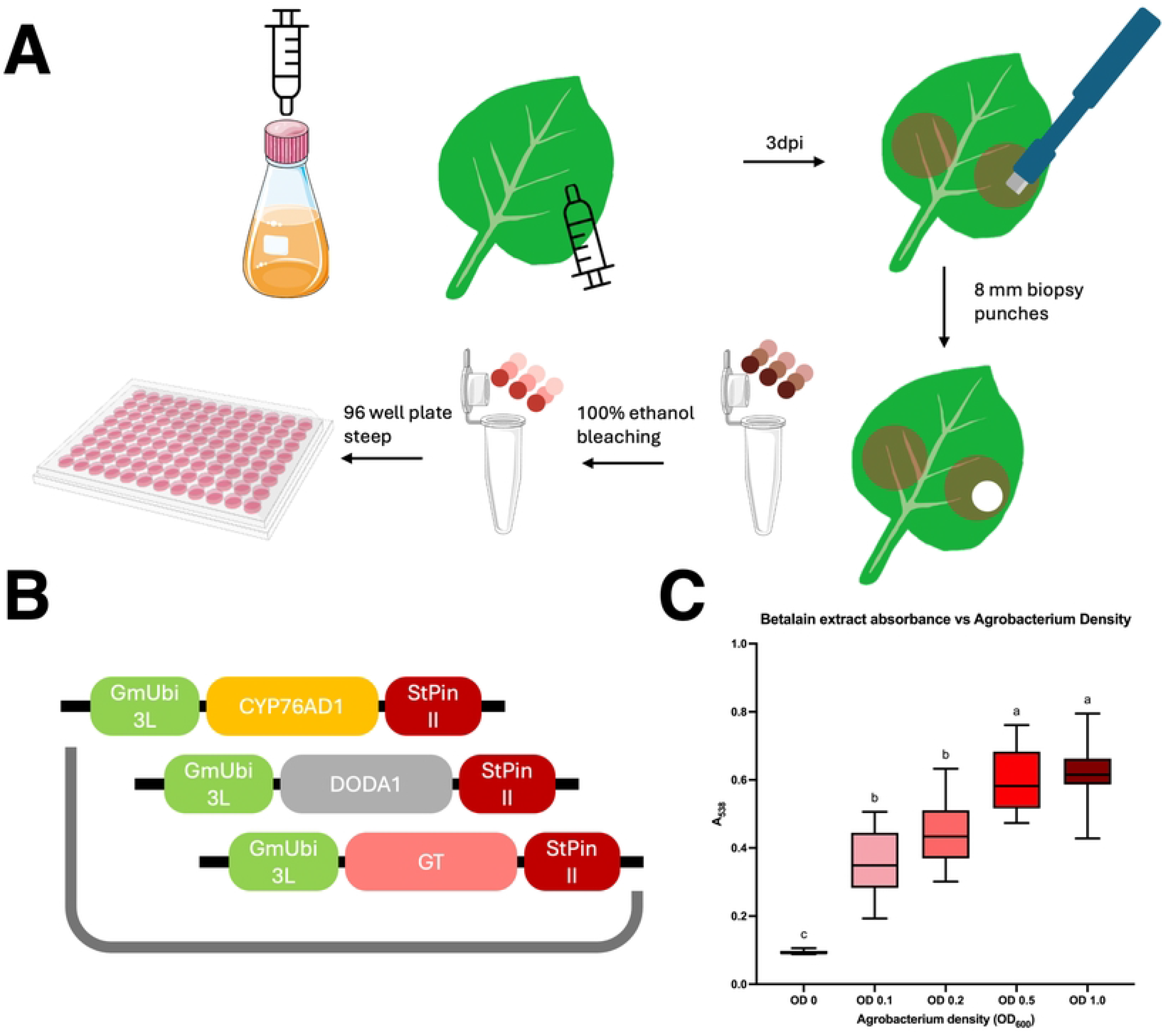
Use of the deconstructed RUBY reporter re-assembled via MultiGreen to confirm vector functionality. A) Workflow for infiltrating and extracting betanin pigments from *Nicotiana benthamiana*. 3-4 week old *Nicotiana benthamiana* cv. TW17 plants were infiltrated with *Agrobacterium* suspension at OD600 ranging from 0, 0.1, 0.2, 0.5, 1.0. Single leaf punches were removed from the infiltration sites. One punch per leaf, three leaves per plant, in three biological replicates performed on different days. Leaf discs were cleared of chlorophyll by soaking in 100% ethanol overnight. Betanin were bled out of the punches by transferring to a 96 well plate containing type I water prior to measuring absorbance at A538 on a Biotek Synergy 2 plate reader. B) Deconstructed RUBY plasmid made using MultiGreen 2.0 stacking the three transcripts of RUBY in tandem. C) Absorbance values for all leaf discs collected. Compact letter display of Tukey HSD comparisons across OD. microtube-open-translucent icon by Servier https://smart.servier.com/ is licensed under CC-BY 3.0 Unported https://creativecommons.org/licenses/by/3.0/.

## Conclusion

MultiGreen expands the original GreenGate cloning architecture by enabling a rapid and efficient means of multiplexing. MultiGreen 1.0 and MultiGreen 2.0 enable multiplexing in series and parallel using components that strictly adhere to the architecture established in the original GreenGate kit. MultiGreen 1.0 takes inspiration from the original GreenGate approach to multiplexing in series, adding the optional H overhang through two new level 0 entry vectors. Alternating the multiplexer modules in MultiGreen 1.0 assemblies rapidly screens out carryover from one concatenation of level 0 modules to the next. Furthermore, MultiGreen 1.0 incorporates the ability to add transcription blockers or spacer sequences between transcriptional units to help minimize interference between successive gene cassettes.

MultiGreen 2.0 introduces multiplexing in parallel within the confines of the default GreenGate assembly architecture to further boost speed of assembly. Parallelized assembly also enables easier exchange of cassettes as needs for a particular gene stack change. In addition, by incorporating two complements of level 1 acceptor vectors for assembly with different antibiotic resistances, and reversion to level 0 modules, transcriptional units can be iteratively stacked in multiples of six. An infinite number of cassettes can therefore be stacked with MultiGreen, only limited by the carrying capacity of the host *E. coli* cloning strain.

The MultiGreen kit encompassing all the vectors and linkers presented in this report for cloning in series and in parallel will be made available through Addgene.

## Supporting information

**SI File: Supplementary methods**

**S1 Fig. Evaluation of level 0 promoter modules in level 1 assemblies with eGFP**. A) Quantitative measurement of GFP on a Synergy2 microplate reader collected with a 485/20 nm excitation filter, 510 nm dichroic mirror, and a 516/20nm emission filter. B) Alignment of the four constructs. pVP272 contains the progenitor promoter module of pVP285; pVP284 contains the progenitor promoter module of pVP286. Level 0 promoter modules used in pVP284 and pVP286 remove out of frame ATG codons after the RBS, introducing Met-gly-ser residues within the A-C level 0 module itself. **S1 Table. List of oligos used in this study**

**S2 Table. List of assemblies and level 0 components used in MultiGreen validation experiments**

**S3 Table. Misassemblies of the protoviolaceinate biosynthesis plasmid**

